# A lifespan single-cell atlas of the human developing hippocampus benchmarks familial Alzheimer’s disease brain organoids

**DOI:** 10.64898/2026.09.18.752796

**Authors:** Ekaterina Ivleva, Emil Kriukov, Joseph F. Arboleda-Velasquez, Petr Baranov

## Abstract

Familial Alzheimer’s disease (fAD) is an early-onset form of AD caused by autosomal-dominant variants in APP, PSEN1, or PSEN2, with PSEN1 accounting for most genetically defined cases [1]. The hippocampus is among the earliest and most severely affected brain regions in AD [2,3]. Human induced pluripotent stem cell (iPSC)-derived brain organoids recapitulate key features of early human brain development and provide a tractable model for studying how fAD mutations perturb neurodevelopmental processes [4]. However, their interpretation is complicated by heterogeneous regional identity, variable maturation state, and cell-type composition across protocols [5,6]. Existing single-cell studies of human hippocampus cover prenatal [7] and postnatal [8–10] stages but do not provide a continuous developmental reference. By elevating the atlas approach in utilizing single-cell RNA-sequencing data, we obtain standardized information on the organoid cell class and type composition and maturation states.

Here, we constructed the Human Developing Hippocampus Atlas (HuDeHA), an integrated single-cell reference comprising 658,059 cells spanning post-conceptional week 3 to 15.3 years, and used it to benchmark iPSC-derived brain organoids carrying PSEN1^E280A^ which is associated with fAD in a large Colombian population. Reference-based mapping revealed altered cellular composition in PSEN1^E280A^ organoids, including reduced radial glia and increased neural crest-derived neurons. These changes were accompanied by cross-lineage transcriptional alterations, including broad upregulation of the ventral patterning factor MEIS2 and reduced expression of the Aβ-binding protein transthyretin (TTR) in choroid-plexus and ependymal-associated populations. Reconstructed neuronal-lineage trajectories showed a shift toward mature states in PSEN1^E280A^ organoids. Together, these findings establish HuDeHA as a resource for developmental benchmarking of hippocampus-relevant organoid systems and describe cell-lineage-specific developmental changes in PSEN1^E280A^ organoids that may inform interpretation of early cellular alterations in fAD.

## Introduction

Familial Alzheimer’s disease (fAD) is caused by autosomal-dominant variants in APP, PSEN1, or PSEN2, with PSEN1 representing the most frequent genetic cause of inherited early-onset AD [1]. Pathogenic PSEN1 variants alter γ-secretase-mediated Aβ production, commonly favoring longer, aggregation-prone Aβ species and increasing the Aβ42/Aβ40 ratio [11,12]. How early these mutations begin to perturb cellular programs during human development is not known [13].

Human iPSC-derived brain organoids provide a tractable system for modeling early human neural development in defined genetic backgrounds. These self-organizing three-dimensional cultures recapitulate aspects of early brain patterning, tissue organization, and neuronal maturation [5,14,15], and offer a uniquely human platform to probe the earliest, presymptomatic consequences of fAD variants at stages that are not accessible in postmortem tissue. Prior fAD organoid studies have primarily characterized amyloid processing, tau pathology, and endosomal or synaptic phenotypes [16,17] , but their effects on early developmental patterning and lineage progression remain poorly understood.

Interpreting organoid phenotypes remains challenging because organoid-derived cell states are often immature, heterogeneous, incompletely specified, and shaped by protocol-dependent differences in regional identity and cell-type composition[5]. Marker-based annotation is particularly challenging for transient progenitors, mixed developmental states, and off-target lineages, making it difficult to distinguish genuine genotype-associated effects from differences in developmental composition. Reference-based mapping to developmental single-cell atlases has therefore emerged as a useful strategy for assigning organoid cells to matched in vivo populations and estimating developmental stage [6] .

To address these questions, we constructed the Human Developing Hippocampus Atlas (HuDeHA), an integrated single-cell reference comprising 658,059 cells spanning every post-conceptional week from PCW3 through PCW16, together with additional later fetal and postnatal time points extending into adolescence. Its broad representation of neural and non-neural lineages provides a framework for resolving developmental cell identities, maturation states, and disease-associated alterations in brain organoids.

We used HuDeHA as an in vivo developmental reference to profile iPSC-derived brain organoids carrying PSEN1^E280A^. Mapping organoid cells onto HuDeHA enabled systematic annotation of cell identity, developmental stage, and neuronal maturation across both hippocampus-relevant and off-target populations. This framework was then used to evaluate genotype-associated differences in cellular composition, transcriptional programs, and neuronal-lineage progression. HuDeHA-based benchmarking showed that PSEN1^E280A^ organoids were characterized by reduced radial glia, increased neural crest-derived neurons, cell-class-specific alterations in developmental and ventricular-associated programs, and a modest shift toward later neuronal maturation states.

## Results

### Human Developing Hippocampus Atlas (HuDeHA) resolves major cell classes across pre- and postnatal timeline

We first constructed the Human Developing Hippocampus Atlas (HuDeHA) by integrating 107 batches comprising 658,059 cells from publicly available scRNA-seq datasets (**Fig. 1A; Fig. S1A, B; Table S1**) spanning from early embryonic with the earliest samples of post conceptional week 3 (PCW3) through postnatal adolescence, with the oldest postnatal samples reaching 15.3 years of age.

**Figure 1.**
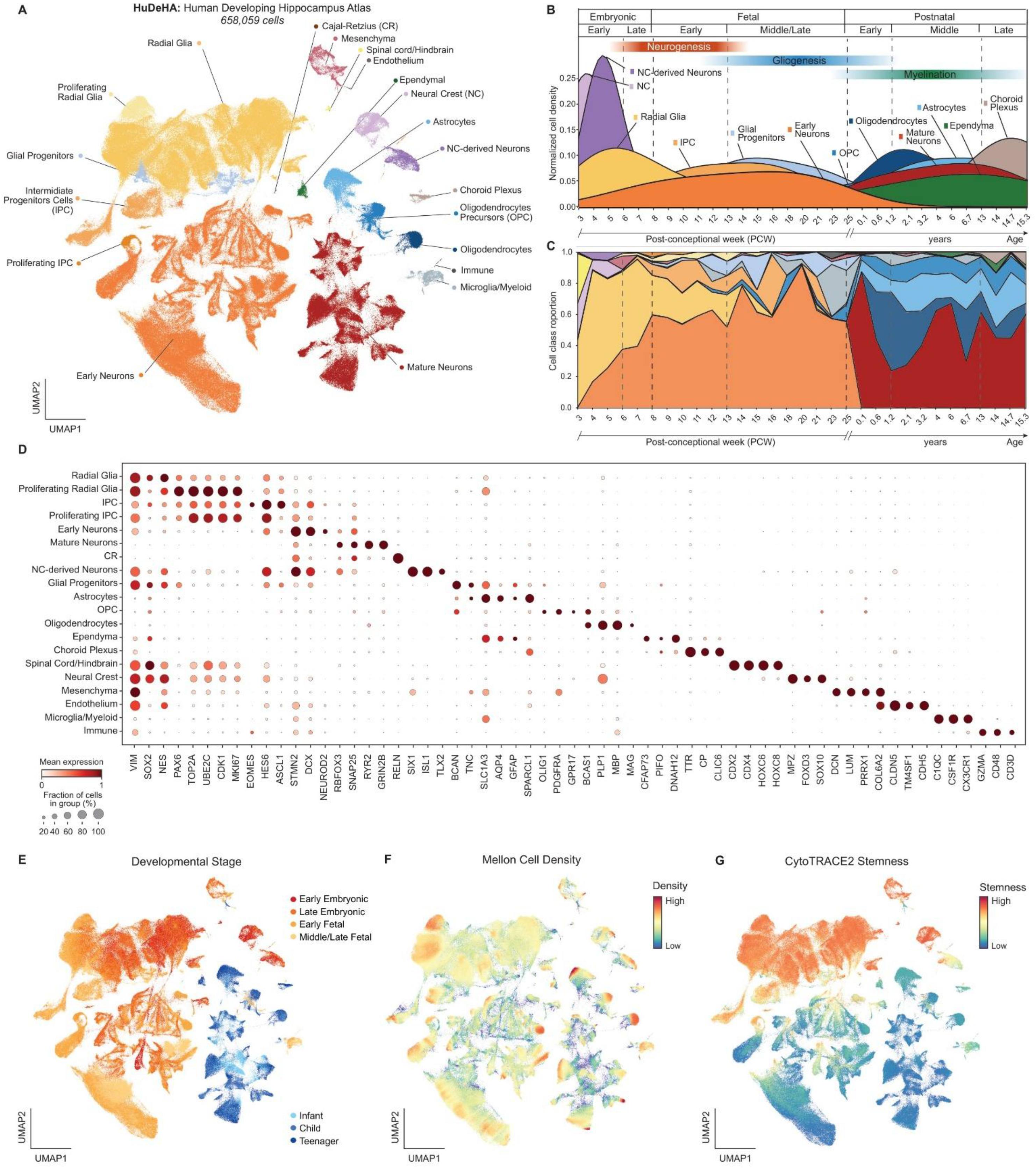
Human Developing Hippocampus Atlas (HuDeHA) captures the temporal dynamics of hippocampal development from early embryonic to postnatal stages. (A) UMAP embedding of the Human Developing Hippocampus Atlas (HuDeHA; 658,059 cells) showing 20 annotated cell classes. The atlas captures a neuronal lineage from radial glia and intermediate progenitor cells (IPCs) to early and mature neurons, together with proliferative radial glia and IPC states; a glial lineage comprising glial progenitors, oligodendrocyte precursor cells (OPCs), oligodendrocytes, and astrocytes; ectodermal non-neuronal populations (ependyma and choroid plexus); neural crest (NC) and NC-derived neurons; non-ectodermal populations (mesenchymal, endothelial, microglial/myeloid, and other immune cells); and a small spinal cord/hindbrain-associated group. (B) Cell class proportion density across development (post conceptional week (PCW) 3 to 15.3 years) showing the expected shifts from neurogenesis to gliogenesis to the emergence of postnatal-associated cell classes: radial glia and NC dominate embryonic samples; IPCs and glial progenitors emerge in fetal stages, while mature neurons together with astrocytes and oligodendrocytes become predominant postnatally. (C) Proportional dynamics of individual cell classes across development, resolved by developmental age. Complementary to (B), this view highlights the timing of individual class emergence and depletion: an embryonic predominance of radial glia and early neurons, progressive loss of NC-derived cells with concomitant expansion of glial progenitors, and postnatal dominance of mature neurons alongside OPCs, oligodendrocytes, and astrocytes. (D) Dotplot showing the expression of canonical markers across all 20 annotated cell classes: radial glia (VIM, SOX2), IPC (EOMES), proliferating radial glia and IPC (TOP2A, MKI67), early (DCX) and mature neurons (SNAP25), Cajal-Retzius (CR) neurons (RELN), glial progenitors (BCAN), OPC (OLIG1), oligodendrocytes (MBP), astrocytes (GFAP), ectodermal non-neuronal populations – ependyma (PIFO), choroid plexus (TTR), NC (MPZ) and NC-derived neurons (ISL1), mesenchyma (DCN), endothelium (CLDN5), microglia (CX3CR1) and immune (CD3D), cord/hindbrain group (CDX2/4). UMAP embedding of HuDeHA color-coded with: (E) Metadata-provided age binned into developmental stage groups, showing age distribution across the embedding. (F) Mellon cell-density overlay, where high-density regions indicate transcriptional states with high concentration of cells (G) CytoTRACE2 potency score, showing the decreasing gradient of stemness from progenitor and precursor populations (top and top left) to differentiated states (bottom and bottom right).

The annotated populations captured the expected diversity of developmental lineages, including neuronal, glial, non-neuronal, vascular, and immune cell types. Using canonical marker genes, we resolved 20 major cell classes in HuDeHA (**Fig. 1D**). Neuroectodermal lineages spanned a neuronal differentiation continuum, beginning with radial glia (194,105 cells) expressing VIM, SOX2, and NES, and proliferating radial glia (11,390 cells) enriched for TOP2A, MKI67, UBE2C, and CDK1. These gave way to intermediate progenitor cells (IPCs; 15,033 cells) marked by EOMES and ASCL1, and proliferating IPCs (3,860 cells), which in turn generated early neurons (246,627 cells) expressing DCX, NEUROD2, and ELAVL3, and more differentiated neurons (95,335 cells) enriched for the post-mitotic and synaptic genes RBFOX3, SNAP25, and SYT1. A small Cajal-Retzius (CR) neurons (170 cells) were distinguished by high RELN expression. The glial lineage included glial progenitors (8,536 cells) expressing BCAN and TNC, which partially overlapped with radial glia in their expression of VIM and SOX2, consistent with a transitional identity. More mature glial populations formed clearly separated clusters of OPCs (11,483 cells; OLIG1 and PDGFRA), oligodendrocytes (7,908 cells; PLP1 and MBP), and astrocytes (15,701 cells; GFAP and AQP4). Additional neuroectodermal populations included ependymal cells (2,200 cells; PIFO) and choroid plexus cells (2,119 cells; TTR), as well as a neural crest (NC) lineage comprising NC cells (10,036 cells; SOX10 and MPZ) and NC-derived neurons (6,302 cells; ISL1 and SIX1). Non-ectodermal populations included mesoderm-derived mesenchyme (8,052 cells; DCN), endothelium (228 cells; CLDN5), microglia (4,876 cells; CX3CR1), and a smaller population of other immune cells (160 cells; CD3D). We also observed a minor spinal cord/hindbrain-related population (480 cells; CDX2 and CDX4) (**Fig. 1D**).

We next examined how cellular composition and maturation state varied across the full atlas timeline, from PCW3 to 15.3 years by visualizing both cell-class proportional density and overall cell-class proportion dynamics (**Fig. 1B, C; Fig. S1D**). These dynamics showed that early embryonic samples were enriched for radial glia and NC populations, whereas later prenatal stages showed a relative decline in radial glia, expansion of early neuronal populations, and emergence of glial progenitor states, consistent with the transition from early neurogenesis toward gliogenic programs. Postnatal samples were enriched for mature neurons and differentiated glial populations, including astrocytes and oligodendrocyte-lineage cells. NC, NC-derived neurons are observed mostly in early embryonic stages.

Donor age from the sample metadata, binned into developmental stages, was projected onto the embedding to show the distribution of developmental stages across cell populations (**Fig. 1E, Fig. S1D**). Mellon-based cell-density estimation highlighted densely sampled regions corresponding to major cell populations and more sparsely sampled regions corresponding to less abundant or transitional states, such as glial progenitors (**Fig. 1F).** To assess developmental potency directly, we computed CytoTRACE2 potency score (**Fig. 1G; Fig. S1C**). Along the neuronal lineage, developmental potency progressively declined from proliferating progenitors and radial glia (median CytoTRACE2 scores = 0.57 and 0.57, respectively), through IPCs (0.43), to early and mature neurons (0.09 and 0.05, respectively), consistent with progressive loss of developmental potential during neuronal differentiation. Other progenitor-like populations, including NC, mesenchyme, and spinal cord/hindbrain-related cells, exhibited high potency scores. In contrast, differentiated postnatal-associated populations, including oligodendrocytes, astrocytes, and ependymal cells, displayed the lowest potency scores.

### HuDeHA allows reference-based mapping of fAD organoids data

The induced pluripotent stem cell (iPSC)-derived brain organoid dataset was obtained from Perez-Corredor et al [18] and included four genotype groups defined by PSEN1 and APOE status: PSEN1^WT^/APOE^WT^, PSEN1 ^WT^/APOE^Christchurch^ (APOE^Ch^), PSEN1^E280A^/APOE^WT^, and PSEN1^E280A^/APOE^Ch^ The dataset comprised eight scRNA-seq libraries generated from eight CRISPR-derived cell lines originating from two donor iPSC backgrounds, with six organoids pooled per cell line prior to sequencing.

We applied scPoli [19], a semi-supervised neural network-based reference-mapping framework that learns a shared latent representation between reference and query cells and transfers reference-derived cell-class labels to the query dataset (**Fig. 2A**). The model was trained on the HuDeHA reference and subsequently used to annotate cells in the brain organoids. This approach resolved progenitor, neuronal, epithelial/ventricular, mesenchymal and NC-derived lineages (**Fig. 2B**). No differentiated glial or immune/vascular populations were detected at this stage.

**Figure 2.**
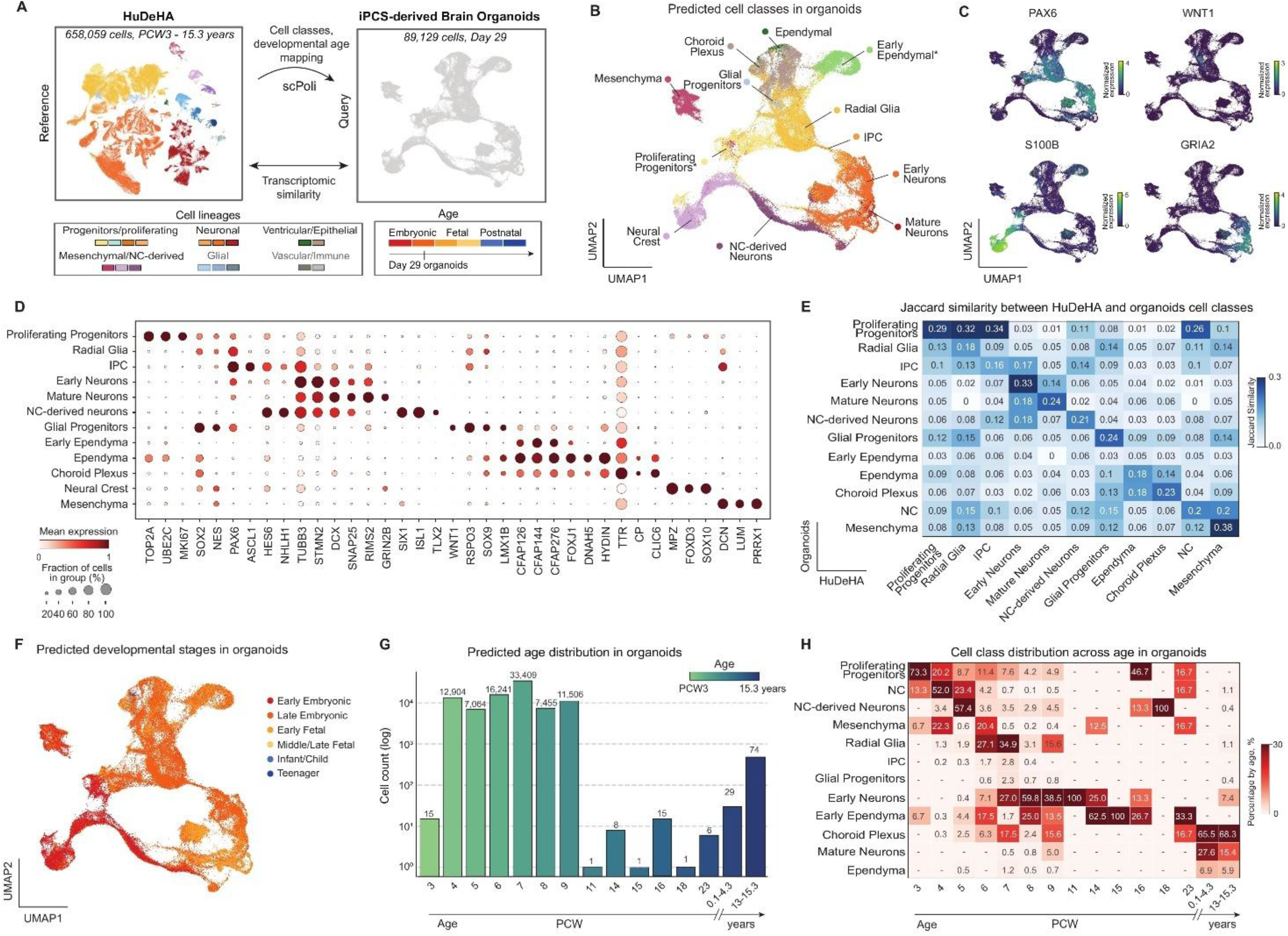
Reference mapping to HuDeHA enables comparison of iPSC-derived brain organoids with in vivo hippocampal cell states and developmental time. **(A)** Schematic of the analysis workflow. HuDeHA (PCW 3 to 15.3 years; 20 annotated cell classes) served as reference for scPoli-based transfer of cell-class and developmental-age labels to iPSC-derived PSEN1^WT^ and PSEN1^E280A^ human brain organoids. scPoli was trained on the intersection gene set between reference and query (22,467 genes), and organoid cells were projected into the learned latent space for label prediction. **(B)** UMAP of iPSC-derived human brain organoids colored by 12 predicted cell classes inferred from HuDeHA, followed by limited manual correction (see Methods). Asterisks mark newly defined or aggregated classes relative to HuDeHA: “early ependyma” (manually added, absent from HuDeHA) and “proliferating progenitors” (aggregate of proliferating radial glia and proliferating IPCs). **(C)** Feature plots of selected regional and lineage markers across the organoid single-cell embedding. PAX6 marked a broad dorsal progenitor compartment, GRIA2 a distinct mature neuronal population, S100B predominantly the NC population with additional expression in ependymal-like cells, and WNT1 a small dorsal midline-like population consistent with roof plate or cortical hem-associated identity. **(D)** Dot plot of marker-gene expression for predicted organoid cell classes: proliferating progenitors (TOP2A, MKI67); radial glia (SOX2); IPCs (ASCL1); early neurons (TUBB3) and mature neurons (GRIN2B); NC-derived neurons (ISL1); glial progenitors (SOX9); early ependyma (CFAP144) and ependyma (FOXJ1); choroid plexus (TTR); NC (MPZ, FOXD3); and mesenchyma (DCN, PRRX1). **(E)** Heatmap of Jaccard similarity between HuDeHA and organoid cell classes, computed on the top 1,500 differentially expressed (DE) genes per class, quantifying overlap between class-specific marker-gene programs. **(F)** UMAP of organoids colored by inferred age bins aggregated into developmental stages. **(G)** Distribution of predicted developmental age on a log-scaled cell-count axis. Cells span broad developmental heterogeneity at a single organoid harvest time point: most cells map to PCW4-PCW9, with a smaller fraction assigned to postnatal stages. **(H)** Heatmap of cell-class proportions across predicted age bins. Early-stage-associated classes (proliferating progenitors, NC-derived neurons) were enriched for younger prenatal ages, while later-stage-associated classes (choroid plexus, ependyma, mature neurons, oligodendrocytes) were enriched for older postnatal ages.

Within the neuronal lineage, we identified IPCs as well as early and mature neurons, although these represented markedly different proportions: early neurons (19,131 cells) and radial glia (18,384 cells) predominated, whereas IPCs (1,308 cells) and mature neurons (877 cells) represented only minor populations. The IPC population expressed ASCL1 and HES6, consistent with an early neurogenic intermediate state, whereas EOMES was detected in <1% of cells. Early neurons robustly expressed TUBB3, STMN2, and DCX, whereas mature neurons expressed GRIN2B and GRIA2 (**Fig. 2C, D**). A strong S100B signal was observed and mapped predominantly to the NC compartment (**Fig. 2B, C)**, which also expressed MPZ, FOXD3, and SOX10 (**Fig. 2D**) with additional mapping to ependymal cells. Although the model showed low annotation-uncertainty scores overall, some populations including ependyma, glial progenitors, and choroid plexus exhibited comparatively higher uncertainty (**Fig. S2A**).

We examined canonical markers together with an alternative embedding of the organoid dataset. The analysis identified a distinct CFAP-enriched population expressing CFAP126, CFAP144, CFAP276, CFAP141, and CFAP90 that lacked a clear counterpart in HuDeHA (**Fig. S2B**). We therefore added this population as an organoid-specific class and annotated it as early ependymal [20]. These cells expressed FOXJ1 and a partial ciliogenesis program but lacked several core axonemal dynein and cilia-assembly genes [6,21]. Ependymal-like cells also showed partial overlap with choroid plexus markers, consistent with incomplete ventricular epithelial differentiation.

The glial progenitor population showed enrichment for roof plate-like/dorsal midline progenitor markers, including LMX1A/B, MSX1, GDF7 (**Fig. S2C**) [22–24]. WNT-family gene expression further highlighted selective WNT1/3A positivity within this population (**Fig. 2C, Fig. S2C**). In contrast, several canonical in vivo glial progenitor markers, including TNC, BCAN, and NFIB, were weakly expressed or absent in organoids (**Fig. S2D**).

Radial glia also showed reduced prediction confidence (**Fig. S2A**), likely reflecting heterogeneous radial glial states in organoids; these cells expressed core radial glia markers such as SOX2 and NES, while also showing partial enrichment for glial progenitor-associated genes.

To evaluate similarity between in vivo and organoid cell classes independently of scPoli-based annotation, we calculated two complementary measures: Jaccard similarity and Pearson correlation [25,26]. Jaccard similarity, computed using the top 1,500 differentially expressed genes (DEGs) for each cell class, quantified overlap between class-specific marker-gene programs in HuDeHA and brain organoids (**Fig. 2E, Fig. S3A-C**). The strongest concordance was observed for mesenchymal cells, which showed the highest matched similarity between organoids and the reference atlas (Jaccard = 0.38). Early neurons also showed clear transcriptional overlap with their corresponding reference population (Jaccard = 0.32), supporting conservation of major neuronal identity in organoids. Several progenitor-related populations showed broader cross-class similarity rather than strict one-to-one matching. For example, proliferating progenitors showed overlap with radial glia and proliferating progenitors in organoids (Jaccard = 0.32 and 0.29, respectively), consistent with shared proliferative and neurogenic marker programs. The early ependymal population showed uniformly low similarity to all reference classes, supporting its classification as an organoid-enriched or atypical population not well represented in the developing hippocampal atlas. As a specificity control, we included reference populations not predicted in organoids, including astrocytes, OPCs, oligodendrocytes, and microglia, and observed no strong similarity to any organoid population (**Fig. S3B**).

Complementary Pearson, Spearman, and Kendall concordance-based analyses across different gene sets supported the same pattern (**Fig. S3D-F**). Mesenchymal cells and early neurons showed the strongest transcriptional similarity to the reference atlas, whereas ependymal, choroid plexus, and early ependymal populations showed lower or more variable similarity, consistent with partial divergence of these organoid states from the in vivo reference.

Using HuDeHA-derived developmental age annotations, we next applied scPoli to predict developmental-stage similarity in the fAD organoid dataset. Predicted organoid states spanned the HuDeHA reference range, from early prenatal to postnatal stages, and showed a modest age-associated organization across the embedding (**Fig. 2F**). Most organoid cells mapped to early prenatal stages, particularly PCW4–PCW9, whereas only approximately 1% mapped to PCW11 or later developmental stages, including postnatal ages (**Fig. 2G**).

We calculated the fraction of cells assigned to each age group within each annotated cell class and visualized these patterns as a heatmap (**Fig. 2H**). The resulting distribution was consistent with expected developmental organization: early predicted ages were enriched among progenitor-like and non-neuronal developmental populations, including proliferating progenitors, mesenchymal cells, and NC cells, whereas later and postnatal-like predictions were concentrated in more differentiated populations, including mature neurons and ependymal-like cells.

### PSEN1^E280A^ organoids display differences in cell-class composition and transcriptional profiles

We next investigated whether PSEN1^E280A^ organoids display alterations in major cell-class composition compared with PSEN1^WT^ organoids (**Fig. 3A-B**). We quantified per-organoid cell-class proportions and assessed statistical support using scCODA, a Bayesian framework for differential compositional analysis of single-cell count data [27] (**Fig. 3B**). This analysis revealed a credible increase in NC-derived neurons and a reduction in radial glia in mutant organoids relative to WT, indicating altered cell-state composition.

**Figure 3.**
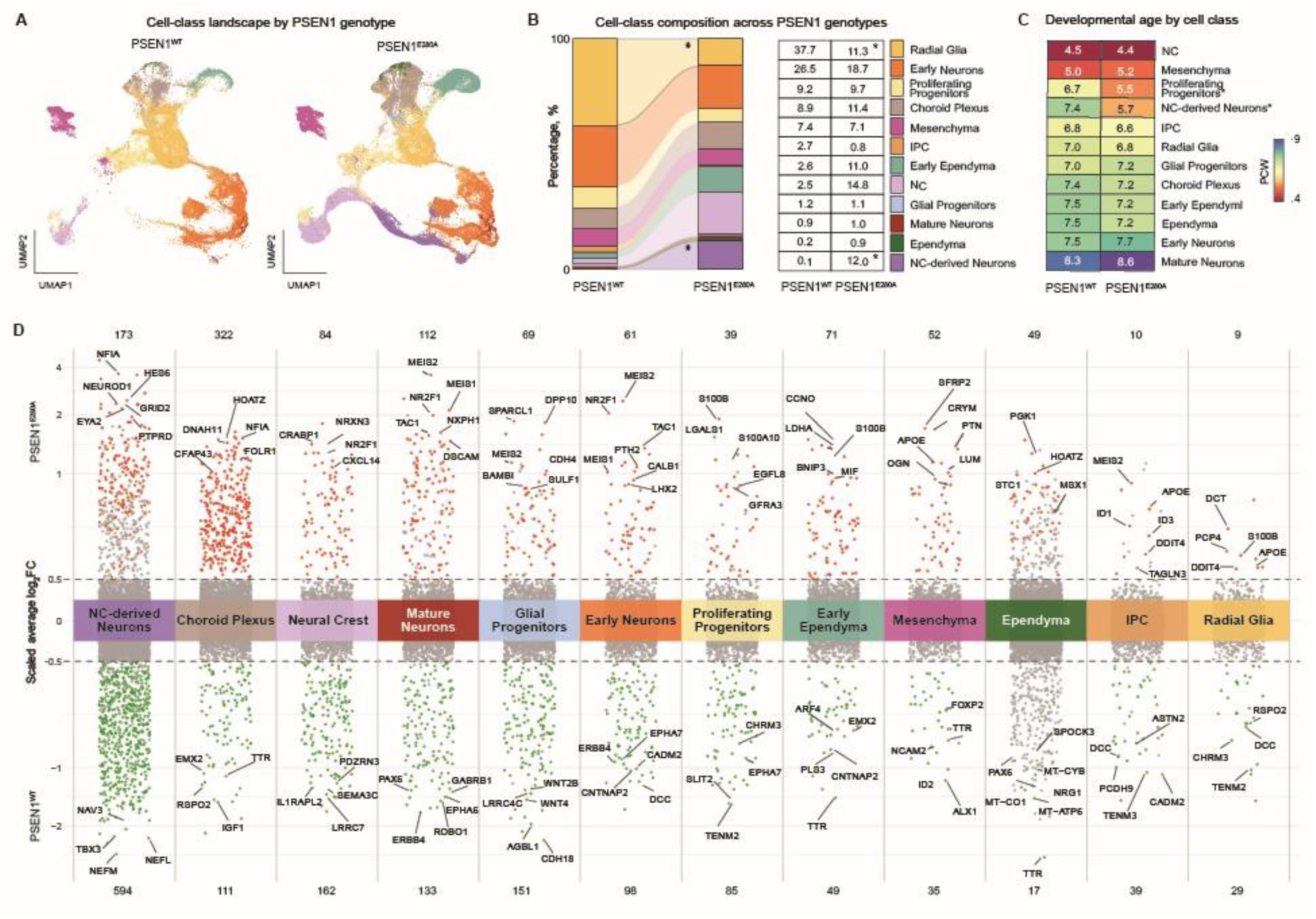
PSEN1^E280A^ is associated with altered lineage balance and cell-class-specific dysregulation of neuronal specification, progenitor, and ventricular programs. **(A)** UMAP of brain organoids split by condition (n = 2 for PSEN1^WT^ and n = 6 for PSEN1^E280A^ organoids), colored by predicted cell class, showing differences in cell-class distribution between conditions. **(B)** Stacked bar plots showing the percentage composition of each cell class by condition. Asterisks indicate cell classes with significantly different relative abundances between conditions. Compared with WT organoids, PSEN1^E280A^ organoids showed an increased proportion of NC-derived neurons and a reduced proportion of radial glia. The table on the right provides the exact percentage of each cell class in each condition. **(C)** Heatmap of the average predicted developmental age by cell class ordered by descending average across the pooled dataset. Asterisks mark cell classes with significant age differences between conditions, assessed using Cliff’s delta. Cutoffs: adjusted p-value ≤ 0.05. **(D)** Differentially expressed genes in PSEN1 ^E280A^ vs ^WT^ organoids, resolved by cell class. Upregulated genes are shown in red and downregulated genes in green; cell classes are ordered by descending total DE-gene count. The top 10 most informative genes per class are labeled; log_2_FC axis is scaled for visualization. Cutoffs: adjusted p-value ≤ 0.05, |log_2_FC| ≥ 0.5.

We next examined predicted age within each annotated cell class and condition (**Fig. 3C**). The resulting heatmap recapitulated the expected developmental ordering, with earlier predicted ages assigned to NC cells, mesenchymal cells, and proliferating progenitors, and later predicted ages assigned to more differentiated populations, including choroid plexus and mature neurons. To quantify condition-associated differences within each cell class, we calculated Cliff’s delta for predicted age distributions. This analysis indicated younger predicted ages for proliferating progenitors and NC-derived neurons in PSEN1^E280A^ organoids compared with WT organoids (5.5 vs. 6.7 PCW and 5.7 vs. 7.4 PCW, respectively), whereas most other cell classes showed negligible differences.

Differential expression analysis within each annotated cell class revealed substantial variation in the number of DEGs, with the highest totals observed in NC and NC-derived neurons, and choroid plexus (**Fig. 3D**).

Among the recurrent transcriptional changes, MEIS2 was upregulated in mature neurons, early neurons, IPCs, and glial progenitors, while MEIS1 was most prominent in mature neurons. NR2F1 was increased in mature neurons, early neurons, and neural crest cells, and TAC1 was elevated in mature and early neurons. In NC-derived neurons, HES6 and NEUROD1 were strongly upregulated together with NFIA, highlighting a distinct neurogenic regulatory program in this population. Progenitor populations showed distinct transcriptional changes. Proliferating progenitors displayed increased S100B, LGALS1, and S100A10, together with reduced SLIT2 and TENM2. Radial glia showed increased S100B and APOE and reduced RSPO2, whereas glial progenitors showed reduced WNT2B and WNT4. IPCs showed increased MEIS2, ID1, ID3, and DDIT4, together with reduced DCC, TENM2, TENM3, PCDH9, and CADM2. Additional changes in adhesion-and axon-guidance-associated genes, including CNTNAP2, ROBO1, and EPH-family genes, were observed across several neuronal and progenitor populations. Ventricular-associated populations showed recurrent reduction of TTR in choroid plexus, early ependymal, and ependymal populations. Choroid plexus cells also showed reduced IGF1 and RSPO2 and increased FOLR1. Early ependymal cells showed increased CCNO, whereas ependymal cells showed increased MSX1 and coordinated downregulation of mitochondrially encoded transcripts, including MT-CO1, MT-CO2, MT-CO3, MT-CYB, and MT-ATP6. The complete list of DEGs is provided in **Table S2.**

We next compared pathway enrichment scores between PSEN1^E280A^ and ^WT^ organoids within each annotated cell class. Pathway enrichment differences were cell-class-specific and most pronounced in proliferating progenitors, where multiple WNT-associated terms were reduced in PSEN1^E280A^ organoids (**Fig. S4A**). To place these pathway shifts in a developmental context, we additionally compared WT and PSEN1^E280A^ organoids with HuDeHA reference cells from matched predicted developmental stages (**Fig. S4B**). This analysis showed that pathway enrichment distributions were broadly overlapping across many differentiated neuronal populations, whereas proliferating progenitors showed the clearest divergence, with PSEN1^E280A^ organoids displaying lower WNT-associated scores relative to WT organoids and age-matched HuDeHA reference cells.

### Neuronal trajectory analysis reveals altered maturation dynamics

We then focused on the neuronal lineage and, using HuDeHA as an in vivo reference, inferred pseudotime across IPCs, early neurons, and mature neurons and visualized the resulting trajectory using a force-directed embedding (**Fig. 4A, B**).

**Figure 4.**
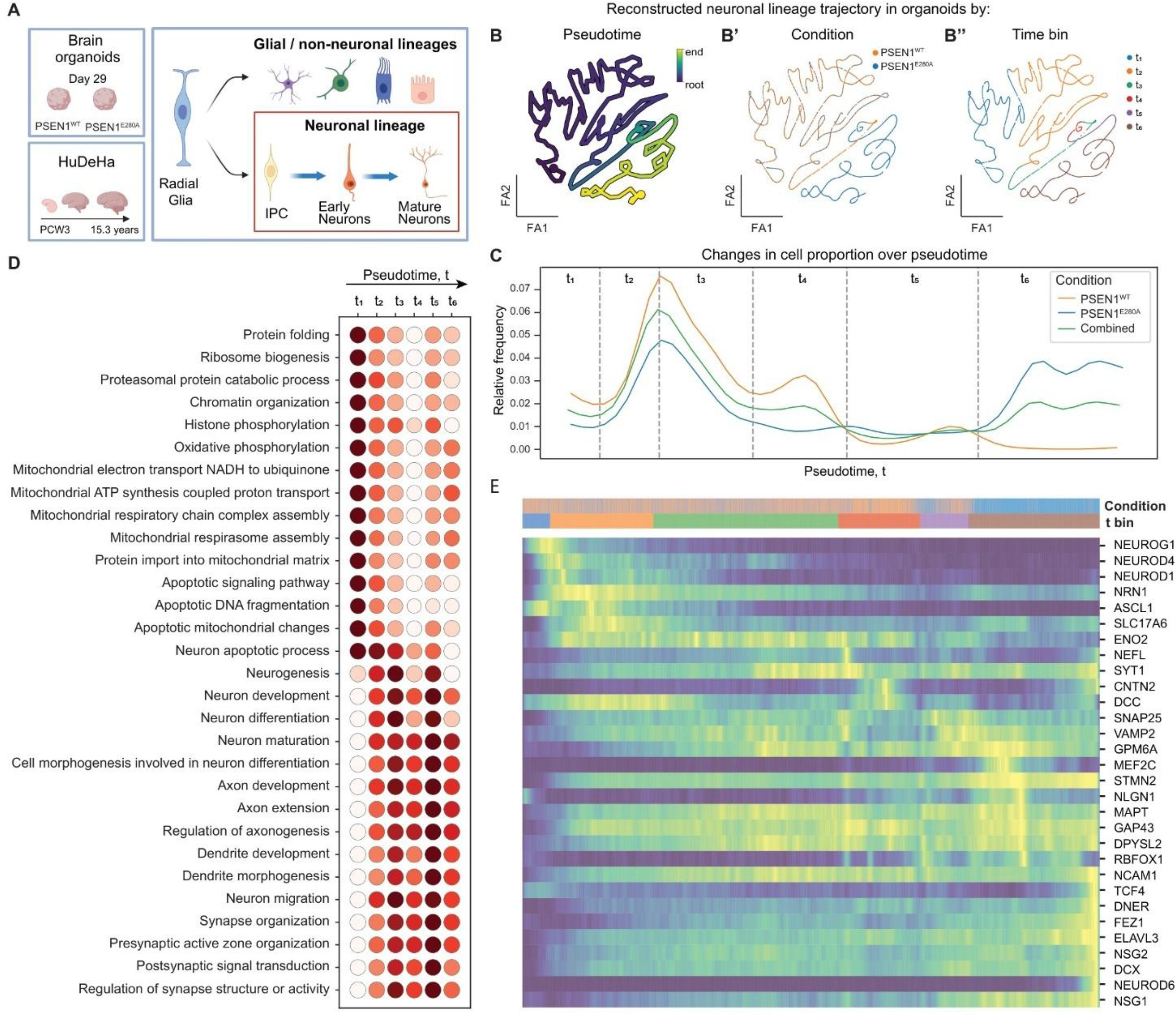
PSEN1^E280A^ alters neuronal differentiation toward more mature developmental states. **(A)** Schematic of the neuronal-lineage trajectory analysis. IPCs, early neurons, and mature neurons were extracted from both iPSC-derived brain organoids and HuDeHA, and used to reconstruct pseudotime trajectories. Organoid trajectories were compared between conditions (PSEN1^WT^ vs PSEN1^E280A^) and benchmarked against the in vivo HuDeHA reference. Reconstructed neuronal lineage trajectory embedding in brain organoids with cells color-coded by: **(B)** inferred scFates pseudotime, representing a continuous developmental gradient. **(B’)** condition (PSEN1^WT^ vs PSEN1^E280A^) **(B’’)** pseudotime bins used for downstream analysis (6 bins) **(C)** Cell-density distribution across pseudotime; major shifts in cell density were used to partition the trajectory into 6 bins. A distinct enrichment at later pseudotime is observed in PSEN1^E280A^ organoids. **(D)** Pathway enrichment scores for neuronal maturation and stress/apoptosis-related processes across pseudotime bins. The data indicate that the late-pseudotime shift in mutant cells is associated with advanced maturation markers rather than elevated cell death or cellular stress. **(E)** Heatmap showing the top 30 genes with the strongest expression changes across organoid neuronal pseudotime. The dynamic gene set includes markers of early neurogenesis and migration (NEUROG1, CXCR4), neuronal differentiation (NEUROD4), and neuronal maturation/cytoskeletal remodeling (SNAP25, SYT1, VAMP2).

The pseudotime ordering was consistent with progressive neuronal differentiation. Proneural transcription factors associated with neurogenic commitment, including NEUROG1, NEUROD1, NEUROD4, and ASCL1, were most prominent at early pseudotime. NEUROG1 and ASCL1 promote neuronal lineage specification, whereas NEUROD1 and NEUROD4 support the transition from progenitor states toward differentiating neurons. In contrast, genes associated with neuronal structure and synaptic function increased toward later pseudotime. These included MAPT, which contributes to axonal microtubule organization; SNAP25, SYT1, and VAMP2, which encode core components of the presynaptic vesicle-release machinery; and NLGN1, a postsynaptic adhesion molecule involved in synapse formation. Together, these expression dynamics indicate a progression from early neurogenic specification toward more differentiated neuronal states with increasing synaptic competence (**Fig. 4E, Fig. S5**).

We next examined the distribution of each genotype along pseudotime and divided the trajectory into six bins corresponding to major shifts in cell abundance (**Fig. 4C, 4B’’**). At the level of biological processes, early bins were enriched for biosynthetic and metabolic programs such as protein folding, ribosome biogenesis, oxidative phosphorylation and mitochondrial respiration, while later bins were progressively enriched for neuron differentiation, neuron maturation, axon and dendrite development, and synapse organization (**Fig. 4D**). Notably, apoptosis-related terms were concentrated at early rather than terminal pseudotime, indicating that the end of the trajectory corresponds to neuronal maturation rather than cell death.

Comparing genotypes along this shared trajectory, PSEN1^WT^ cells were concentrated at earlier-to-intermediate pseudotime, peaking around t2–t3, whereas PSEN1^E280A^ cells were comparatively more represented at later pseudotime bins (**Fig. 4B′, C**). This modest redistribution indicates a relative enrichment of the mutant population for more mature neuronal-lineage states, without an accompanying change in the overall structure of the trajectory. Together, these analyses show that PSEN1^WT^ and PSEN1^E280A^ organoids traverse the same neuronal-lineage trajectory but differ modestly in their distribution along it, with the fAD mutation associated with a shift toward later, more mature pseudotime states.

We extracted the corresponding neuronal-lineage populations from HuDeHA and reconstructed an in vivo trajectory using the same analytical framework (**Fig. S6A**). The distribution of annotated HuDeHA cell classes and heatmap of metadata-defined developmental stages along pseudotime followed the expected developmental order (**Fig. S6B, D**). Relative enrichment of cells in the latest pseudotime bin was not detected in HuDeHA (**Fig. S6C**), indicating that the relative enrichment of late-pseudotime states in mutant organoids is not simply a general property of the reference trajectory. We divided HuDeHA neuronal-lineage cells into five pseudotime bins defined by major changes in cell abundance along pseudotime (five bins for HuDeHA, versus six for the organoid trajectory. We assessed the similarity between organoid and HuDeHA neuronal trajectories using Pearson correlation (**Fig. S6E**). Across all genotype groups, organoid neuronal states were most similar to the early HuDeHA reference bins, particularly t1–t2, with progressively lower similarity to later reference bins, indicating that all organoid genotypes remain predominantly early-like relative to the in vivo developmental trajectory. However, PSEN1^E280A^ organoids showed modestly higher similarity to the intermediate HuDeHA t3 bin at later organoid pseudotime than WT organoids.

## Discussion

We established the Human Developing Hippocampus Atlas (HuDeHA) reference spanning PCW3 to 15.3 years. Its broad temporal coverage captures major transitions from early embryonic neurogenesis through prenatal hippocampal development and postnatal maturation. The existing hippocampal references are restricted to a relatively narrow age window: atlases of the developing hippocampus cover fetal stages [7], whereas others capture only postnatal glial diversity [10] and integrating data across such disparate ages is nontrivial. To extend coverage of the earliest stages, we incorporated early prenatal data from broader brain-region studies and filtered for hippocampus-related cells, thereby broadening the developmental range and increasing the number of sampled cells. By integrating multiple datasets into a common reference, HuDeHA provides extensive representation of neuronal, progenitor, glial, vascular, immune, and ventricular/epithelial populations across developmental stages. This enables systematic comparison of cell identities and maturation states, supports the annotation of in vitro models against their in vivo counterparts, and provides a framework for identifying developmental populations or trajectories that may be altered in disease-associated organoids.

Using HuDeHA, we found that the fAD iPSC-derived brain organoids carrying the PSEN1^E280A^ mutation predominantly recapitulated early embryonic development, with most cells mapping to approximately PCW 5-9. This early first-trimester identity is consistent with the progenitor-rich composition of unguided cerebral organoids at this differentiation stage and their limited similarity to later fetal reference states [6] .

### Reference mapping identifies divergent and organoid-specific populations

HuDeHA-based annotation showed that brain organoids contain a prominent neuronal lineage together with neural crest-like and mesenchymal compartments, whereas differentiated glial populations were not detected at this stage. Although neural crest cells may appear unexpected in a cerebral organoid workflow, previous single-cell studies have reported mesenchymal, neural crest, and other off-target non-neural cell classes across brain organoid protocols [6,14]. The neural crest like compartment observed here may therefore reflect incomplete regional restriction or protocol-specific expansion of non-neural states, rather than being itself a mutation-driven phenotype.

Within the neuronal lineage, radial glia and early neurons predominated, whereas IPCs and mature neurons represented only minor populations, likely reflecting the developmental stage and differentiation characteristics of this organoid model. Chimeric transcriptional profiles of organoid cells are a further challenge for annotation and understanding the transcriptional similarity with in vivo reference is critical. Thus, choroid plexus- and ependymal-like populations showed limited similarity to their corresponding HuDeHA reference states, consistent with incomplete maturation of ventricular cell classes in brain organoids [4,5]. We also identified an abundant immature ventricular epithelial population with partial activation of the ciliogenesis program but no clear counterpart in HuDeHA. Its modest FOXJ1 expression and absence of several core axonemal and dynein-assembly genes suggest an organoid-specific early ciliogenic state rather than fully mature multiciliated ependyma. Glial progenitor-like cells in the organoids also showed limited similarity to their in vivo counterparts. Closer examination revealed expression of multiple roof plate and cortical hem markers, suggesting that this population may instead represent a cortical hem-like or choroid plexus-adjacent signaling state.

### PSEN1E280A is associated with altered lineage allocation and cell-class specific transcriptional programs

Compositional analysis suggests that PSEN1^E280A^ is associated with altered lineage balance in organoids, including a relative reduction in radial glia and an increase in neural crest populations. This imbalance may reflect altered progenitor behavior in PSEN1^E280A^ organoids, including reduced radial glia maintenance, increased neuronal differentiation. Because neural crest abundance can also vary with the differentiation protocol, the increase in neural crest populations is best interpreted as an association with the mutation rather than a direct consequence of it. Together, these shifts are consistent with the idea that early lineage decisions can be biased when developmental signaling programs are perturbed [28].

Differential expression analysis indicated that PSEN1^E280A^-associated changes extended beyond differences in cell class composition and involved transcriptional alterations across multiple developmental lineages. Three broad patterns emerged: changes in regional and neuronal-fate specification, altered progenitor and neurogenic programs, and disruption of ventricular epithelial and choroid plexus-associated states.

We observed recurrent upregulation of MEIS2 across neuronal lineage including IPC, early and mature neurons, accompanied by increased MEIS1, NR2F1, and TAC1 expression in early and mature neurons. MEIS proteins are TALE-class homeodomain transcription factors involved in tissue- and lineage-specific transcriptional regulation [29], while NR2F1 regulates regional progenitor dynamics and cortical developmental identity [30]. The recurrent MEIS2 signal may reflect a broader alteration in regional patterning or developmental identity in PSEN1^E280A^ organoids. However, the functional significance of this change remains uncertain because MEIS2-associated transcriptional programs are context dependent.

Radial glia in PSEN1^E280A^ organoids showed increased S100B and APOE, suggesting an altered progenitor state rather than a demonstrated shift toward astrocytic differentiation. Because both genes are linked to AD-associated glial responses [31], their dysregulation in an early progenitor population may indicate that disease-relevant transcriptional changes emerge before mature glial populations are established.

The increased expression of HES6 and NEUROD1 in NC-derived neurons suggests that PSEN1^E280A^ affects the regulation of early neurogenic programs within this population. HES6 can relieve HES1-mediated repression and support proneural-factor activity, whereas NEUROD1 promotes neuronal differentiation [32–34]. However, because these cells mapped to younger developmental stages, their co-upregulation is more consistent with persistence or prolongation of an immature differentiating state than with advanced neuronal maturation.

Ventricular and choroid plexus-associated populations showed a distinct set of transcriptional alterations, most notably recurrent reduction of TTR. TTR is a major secreted product of the choroid plexus and has been reported to bind Aβ, limit its aggregation and toxicity, and facilitate its transport across brain barriers [35–37]. Choroid plexus dysfunction and impaired Aβ clearance have also been described in AD models [38]. Reduced TTR may reflect altered maturation or secretory function of ventricular-associated populations and could potentially affect Aβ homeostasis at later stages.

Previous studies have shown that reduced presenilin function can impair Notch signaling and promote premature neuronal differentiation, while familial PSEN1 mutations have also been associated with altered neurogenic timing in human stem-cell and organoid models [39]. In our data, Notch-associated pathway scores were only modestly reduced and primarily in proliferating progenitors. Thus, our findings are broadly consistent with altered progenitor maintenance or developmental timing, but they do not support widespread Notch suppression or establish reduced Notch activity as the direct cause of the observed compositional and maturation changes. The reduction of WNT-associated programs in proliferating progenitors provides a plausible link between altered developmental signaling, the loss of radial glia, and the shift toward more advanced neuronal states. Functional perturbation studies will be required to determine whether these pathway changes directly drive the observed phenotype.

Developmental-age mapping indicated that the organoids predominantly recapitulate embryonic stages, with most cells assigned to approximately PCW 5-9 that placing the organoids within an early first-trimester window. This result is consistent with the early, progenitor-rich composition expected of unguided cultures at this differentiation stage and with their limited similarity to later fetal reference states [6]. Within this early window, PSEN1^E280A^ organoids showed a modest shift toward younger predicted ages in proliferating progenitors and NC-derived neurons relative to PSEN1^WT^.

### Neuronal trajectory analysis uncovers altered developmental progression

Trajectory analysis of the neuronal lineage provided a framework for comparing maturation dynamics across genotypes and for identifying candidate programs associated with altered developmental progression. In our dataset, PSEN1^E280A^ was associated with a shift toward more advanced states along the neuronal maturation trajectory. These later states showed stronger expression of genes involved in neuronal differentiation, axon and dendrite development, and synaptic organization. This observation is broadly consistent with prior studies showing that some fAD mutations can promote premature neuronal differentiation or accelerated progression out of progenitor states [1,13]. At the same time, other studies have reported the opposite pattern, including increased progenitor retention and reduced neuronal differentiation in mutation-specific models such as PSEN1^L435F^ [40]. These contrasting findings suggest that the effect of PSEN1 mutations on neuronal development is likely mutation-, model-, and stage-dependent. Importantly, the shift observed here reflects relative advancement within the organoid developmental trajectory rather than acquisition of fully mature neuronal identity.

Overall, our findings indicate that PSEN1^E280A^ is associated with altered developmental cell-state balance in brain organoids, involving progenitor, neuronal, neural crest-like, and ventricular epithelial-like populations. HuDeHA provided a useful framework for distinguishing hippocampus-relevant neural states from organoid-specific populations and for interpreting mutant-associated changes in the context of human in vivo development. Together, these results support the value of reference-based benchmarking for improving interpretation of organoid models and for placing disease-associated cellular changes within a developmental framework.

### Limitations

As for the limitations of the study: the organoid dataset included only two PSEN1^WT^and six PSEN1^E280A^ organoids from a single published study, limiting statistical power and the ability to distinguish mutation-associated effects from organoid-to-organoid variability; Observed increase in NC-derived populations may partly reflect protocol-dependent lineage specification rather than a direct consequence of PSEN1^E280A^.

## Methods

Data collection and preprocessing for HuDeHA assembly

### Data collection

HuDeHA spans developmental ages from PCW3 to 15.3 years. We organized source public datasets into three groups according to developmental stage and anatomical annotation: early development, prenatal hippocampus, and postnatal hippocampus. Each developmental group was integrated separately. Early-development datasets were filtered to retain brain-, forebrain-, and hippocampus-relevant populations, whereas the prenatal and postnatal groups were restricted to hippocampus-annotated tissues. The retained subsets were subsequently separated by original batch, reprocessed as individual objects, and jointly integrated to construct HuDeHA.

The early development dataset included 70 batches and 476,237 cells derived from three published studies [41–43]. The prenatal dataset included 15 batches and 48,679 cells from three studies [7,42,44]. The postnatal dataset included 22 batches and 133,143 cells from two studies [8,10]. Data sources and Metadata for each batch is provided in **Table S1.**

The early-development group included broader anatomical sources, including whole-body, whole-head, whole-brain, and ganglionic eminence samples. Hippocampus-specific dissections were not consistently available at the earliest developmental stages. Following separate integration, these datasets were conservatively filtered using established developmental, regional, and lineage markers to retain brain-, forebrain-, and hippocampus-relevant populations.

Ganglionic eminence datasets were included to improve the representation of progenitor populations contributing to inhibitory-neuron development. This inclusion was motivated by evidence that hippocampal GABAergic interneurons arise from ganglionic eminence progenitor zones and migrate into the developing hippocampus [45,46]. Consistent with this developmental origin, human single-cell studies have identified distinct medial and caudal ganglionic eminence-associated inhibitory lineages during fetal development [43,47].

This stage-specific dataset-selection strategy enabled the construction of a continuous developmental reference containing hippocampus-relevant cell populations across developmental periods that are not uniformly represented by hippocampus-specific datasets.

### Individual objects processing

Each dataset was processed independently using the Seurat package (v4.3.0) [48–50]. Each sample was treated as a separate Seurat object. Quality control (QC) was applied using the following thresholds: percent.mt < 30%, percent.rb < 40%, nCount_RNA > 300, and nFeature_RNA > 400. Upper thresholds for gene and UMI counts were determined based on dataset-specific distributions. Following QC, each object was normalized, variable features were identified using FindVariableFeatures(selection.method = “vst”, nfeatures = 3000), and data were scaled using vars.to.regress = c(”percent.mt”, “percent.rb”, “S.Score”, “G2M.Score”). Cell cycle scores were computed using the CellCycleScoring() function with the cc.genes.updated.2019 gene set [51].

To remove potential doublets, we applied the DoubletFinder package (v2.0.3) [52]. The discovery ratio parameter (pK) was optimized for each object by sweeping across multiple values using paramSweep_v3 and selecting the value with the highest Binary Classification Metric (BCmetric) using find.pK. The expected doublet rate was set to 5% of the total cell count and adjusted for undetectable homotypic doublets using estimates from modelHomotypic. After doublets removal, each Seurat object was reprocessed.

### Intermediate atlases assembly

#### Early development stage

To filter early hippocampus part for early developmental stage group we split the data into 3 groups: whole body and whole head jointly (PCW3 –8), whole brain (PCW7–12), and ganglionic eminence datasets (PCW7-PCW16) separately. Whole body and whole head samples were integrated together and then filtered to retain only brain-derived cells. Whole brain datasets were integrated separately and used to select hippocampal and adjacent telencephalic populations. Ganglionic eminence datasets were processed as an additional compartment and later included in the combined integration.

#### · Whole body + whole head

Whole-body and whole-head datasets covering PCW3–8 were jointly integrated, and the resulting object was subsequently restricted to brain-derived cells. As an initial strategy, we performed whole-brain reference-based mapping using MapMyCells (RRID:SCR_024672) and identified putative brain cells based on predicted class probabilities, evaluating thresholds of 0.75 and 0.99. While the more stringent threshold (0.99) produced slightly clearer separation of predicted brain and non-brain populations, overall discrimination remained suboptimal. We therefore complemented this strategy with literature-derived early developmental brain markers SOX2, VIM, HES1/HES5, PAX6, NES [53] to refine the brain-specific subset.

#### • Whole brain

Whole-brain datasets spanning PCW7–12 were jointly integrated, after which the dataset was restricted to forebrain and hippocampus-relevant populations based on established marker genes, including ZBTB20, CALB1, FOXG1, PAX6, LHX5, EMX2, OTX2, NKX2-1 [41,42], [53–55].

#### • Ganglionic eminences

All available ganglionic eminence (GE) datasets spanning PCW7-16 were jointly integrated, and differentiation potency was quantified for each cell using CytoTRACE2 [56] with default parameters. We then evaluated a reference-based mapping approach for GE cell annotation using publicly available reference datasets. Because the resulting annotations showed poor concordance with established marker-gene expression and expected regional identity, this reference-mapping approach was not used for final annotation. To enrich for progenitor populations and reduce the contribution of differentiated GE-derived cells, downstream inclusion was restricted to cells classified as multipotent by CytoTRACE2 (CytoTRACE2_Potency == “Multipotent”).

#### Prenatal, postnatal stages

Prenatal and postnatal hippocampal datasets were processed and integrated separately. Contaminating populations, including erythrocytes, were identified by marker expression and removed before final integration.

### HuDeHA integration

Following the intermediate integration of the early developmental, prenatal, and postnatal datasets, hippocampus-enriched subsets were retained. The resulting data were then separated by original batch and reprocessed as individual objects, yielding 107 batches that were subsequently integrated to construct HuDeHA.

For the final integration, an scVI model [57,58] was trained on 3,000 highly variable genes, with the original batch specified as the ‘batch_key’. Hyperparameter tuning evaluated combinations of 128 or 256 hidden units, one to three hidden layers, 20, 30, or 40 latent dimensions, and maximum training durations of 100 or 300 epochs. The final architecture, comprising three hidden layers, 256 hidden units, and 30 latent dimensions, was selected based primarily on the lowest validation loss, supported by inspection of the training and validation loss curves and evaluation using scIB metrics and gene expression distribution across the embedding. The final model was trained for up to 100 epochs with early stopping enabled, using validation loss as the monitored metric and a patience of five epochs.

Cell populations in the scVI latent space were initially annotated using a curated marker-gene list. The trained scVI model was then converted to an scANVI model [59] , using the initial annotations as supervision and retaining the same underlying model architecture. scANVI was used to refine and propagate labels across the complete dataset, including cells that remained unannotated after the initial marker-based annotation.

Brain organoids data processing

### Data preprocess and filtering

The data were deposited from GEO accession **GSE241453**. Each sample was processed using the same pipeline described above, including mitochondrial and ribosomal QC filtering, doublet removal, cell-cycle scoring, and scaling.

After preprocessing, datasets were integrated using default Seurat v.4 integration pipeline, as described above. After initial annotation, we identified one Seurat cluster enriched for mitochondrial genes, consistent with low-quality or stressed cells. These 9,359 cells were removed, and the remaining dataset was reintegrated in Seurat using 30 principal components (PCs). For manual cell-class refinement, we also generated an alternative scVI embedding (n_layers = 2, n_latent = 20, n_hidden = 128) and used this representation to refine the cell-class labels initially transferred by scPoli. Labels were refined by jointly evaluating transferred labels, cluster structure in the scVI latent space, and canonical marker-gene expression (**Fig. S2F-G**).

Cell class mapping

To transfer cell class annotations from the reference atlas to our brain organoid dataset, we used scPoli [19] with the following preprocessing and training procedure. First, we harmonized cell-type granularity in both reference and query by collapsing closely related subtypes: IPC proliferating and Radial glia proliferating were merged as Proliferating progenitors, CR neurons were grouped with Mature neurons, Immune and Microglia were merged as Microglia. To reduce overfitting and ensure a comparable feature space, we performed low-expression gene filtering separately in the reference and query datasets and then restricted the model to the 22,467 genes retained in both datasets. scPoli was trained with an embedding dimensionality of 10, a latent dimensionality of 20, one hidden layer containing 128 units, and a negative-binomial reconstruction loss. Early stopping monitored validation prototype loss in minimization mode with a patience of 20 epochs. The learning rate was reduced on plateau using a patience of 13 epochs and a reduction factor of 0.1. The checkpoint with the lowest validation prototype loss was retained.

Developmental age mapping

To harmonize temporal resolution and reduce class imbalance, developmental ages with limited cell representation were combined into broader bins as follows: PCW23 and PCW25 as PCW23–25; 0.1– 4.3 years; 6–6.7 years; and 13–15.3 years. These merges produced more balanced age-category frequencies in the reference atlas. Because ground-truth developmental ages were unavailable for organoid cells, downstream analyses were restricted to high-confidence predictions, retaining cells with a maximum posterior class probability of at least 0.75.

Developmental-age prediction used the same scPoli model architecture and training parameters described above, with developmental-age group used as the supervised label.

Compositional analysis

Cell-class compositional differences between PSEN1^E280A^ organoids (MUT; n = 6) and wild-type organoids (WT; n = 2) were assessed using scCODA, a Bayesian model designed for compositional single-cell data analysis [27]. scCODA’s automatic reference-selection procedure identified Mature Neurons as the reference cell class. Because this population was relatively low in abundance, robustness to reference choice was assessed by repeating the analysis using each retained cell class as an alternative reference. The inferred effects for radial glia and NC-derived neurons were consistent across reference choices.

Developmental age comparison in brain organoids

To compare predicted developmental ages between PSEN1^E280A^ and PSEN1^WT^ organoids, analyses were performed separately within each cell class. Predicted age values were converted to a continuous numerical scale and compared between genotypes using a two-sided Mann-Whitney U test. Cell classes represented by fewer than 10 cells in either genotype group were excluded from statistical testing. P-values were adjusted for multiple testing across cell classes using the Benjamini-Hochberg false discovery rate (FDR) procedure. Effect sizes were quantified using Cliff’s delta, where positive values indicate older predicted developmental ages in PSEN1^E280A^ cells relative to WT cells. Adjusted p-value 0.05 was used as statistically significant.

Differential expression in HuDeHA and brain organoids

Differential expression analysis between PSEN1^E280A^ and PSEN1^WT^ brain organoids was performed separately within each cell class using Seurat FindMarkers. For each cell class, cells were subsetted by condition and differential expression was tested using the Wilcoxon rank-sum test Differentially expressed genes were defined as those with an absolute log_2_fold change ≥ 0.5 and an adjusted p-value < 0.05.

Pathway enrichment in HuDeHA and brain organoids

For pathway enrichment analysis, we performed ssGSEA using a curated subset of GO and HP gene sets related to Notch and WNT signaling, neuronal and glial differentiation, and cell-fate specification, selected from the MSigDB C5 collection. Per-cell enrichment scores were computed with the escape package v1.10.0 [60] using enrichIt function with default parameters. To compare brain organoids with HuDeHA, reference atlas cells were restricted to the developmental age bins predicted in the organoid dataset and analyzed within matched cell classes. This minimized developmental-stage effects and enabled comparison of pathway activity in a more comparable cellular context.

Jaccard similarity

To quantify transcriptional similarity between HuDeHA and brain organoids across matched cell classes or developmental time points, we compared group-specific marker gene sets using Jaccard similarity. Differential expression was performed independently within each dataset using scanpy.tl.rank_genes_groups with the Wilcoxon rank-sum test. For each cell class or time-point group, genes were ranked by the Scanpy test statistic, and the top *n* genes were selected to define a group-specific marker set. Only genes detected in both datasets were considered for downstream comparison.

For each matched HuDeHA -brain organoids pair, Jaccard similarity was calculated as:

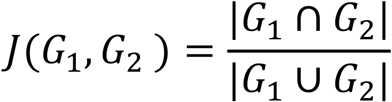

where *G*_1_ and *G*_2_ represent the top-ranked gene sets from the matched HuDeHA and brain organoids groups, respectively.

Transcriptomic similarity between HuDeHA reference cell classes and brain organoid cell classes was assessed using cell-class-level mean expression profiles. Similarity was evaluated using four feature sets: 3,000 highly variable genes (HVGs) identified in HuDeHA, 3,000 HVGs identified in the brain organoid dataset, the intersection of the two HVG sets (944 genes), and their union (5,056 genes). Mean normalized expression was calculated for each gene within each cell class, and pairwise similarity between HuDeHA and brain organoid cell classes was quantified using Pearson, Spearman’s rank, and Kendall’s rank correlations.

Cell fates trajectory inference

To compare neuronal differentiation trajectories between HuDeHA and brain organoids datasets, we subset neuronal-lineage populations from both datasets, including intermediate progenitor cells (IPCs), early neurons, and mature neurons. To reduce imbalance in cell numbers between the reference atlas and organoid datasets, the HuDeHA neuronal-lineage subset was randomly downsampled to 18,000 cells, closely matching the 17,296 neuronal-lineage cells in the brain organoid dataset. Downsampling was performed using a fixed random seed to ensure reproducibility.

Trajectory inference was performed on the scVI-derived latent representation to reduce the influence of technical variation and batch-specific effects.

We first used Palantir [61] to construct a k-nearest-neighbor graph using 30 neighbors and computed diffusion components with n_eigs = 2. Pseudotime inference and trajectory visualization were then performed using scFates [62]. For visualization, a ForceAtlas2 layout was initialized using the first two components of the trajectory representation.

Cell potency and density analysis

Cellular potency across HuDeHA was estimated from single-cell RNA-seq profiles using CytoTRACE2 [56] with default parameters. Cell-state density was estimated using Mellon [63] across all HuDeHA cells in the scANVI latent space, using all 20 latent dimensions (n_components = 20). Mellon generated a log-density estimate for each cell, representing its local abundance within the integrated latent space. All other Mellon parameters were left at their default values.

## Authors contribution

E.I. designed and conducted the computational analyses, performed data integration and downstream bioinformatics analyses, interpreted the results, prepared the figures, and drafted the manuscript. E.K. contributed to the analytical design and methodological framework, supervised the bioinformatics analyses, and critically reviewed the code, figures, results, and manuscript. J.A. generated the publicly available brain organoid dataset analyzed in this study and contributed expertise related to the experimental model. P.B. conceived and supervised the study, provided overall scientific direction, contributed to interpretation of the results, and critically revised the manuscript. All authors reviewed and approved the final manuscript.

## Acknowledgements.

The authors want to thank the research groups who generated the original datasets used in this study. This work was supported by the grants from Gilbert Family Foundation (PB) and MGB GCTI (PB). We also acknowledge funding from the National Institutes of Health (**RM1NS132996-01).**

## Data availability

This paper analyzes existing, publicly available data, referenced and described in **Table S1**. For organoids we used data deposited from GEO accession **GSE241453**. We do not claim any authorship over the original datasets provided in **Table S1** and labeled as not published at the moment of the manuscript submission.

Requests for further information and resources should be directed to and will be fulfilled by the lead contact, Petr Baranov.

Any additional information required to reanalyze the data reported in this paper is available from the lead contact upon request.

## Code availability

All original code has been deposited at GitHub and is publicly available at https://github.com/ivlevae/Alzheimer/tree/main Alzheimer as of the date of publication.

