## Supplementary material for "A lifespan single-cell atlas of the human developing hippocampus benchmarks familial Alzheimer’s disease brain organoids": Figures and legends

Supplementary Materials

**Supplementary Figures**

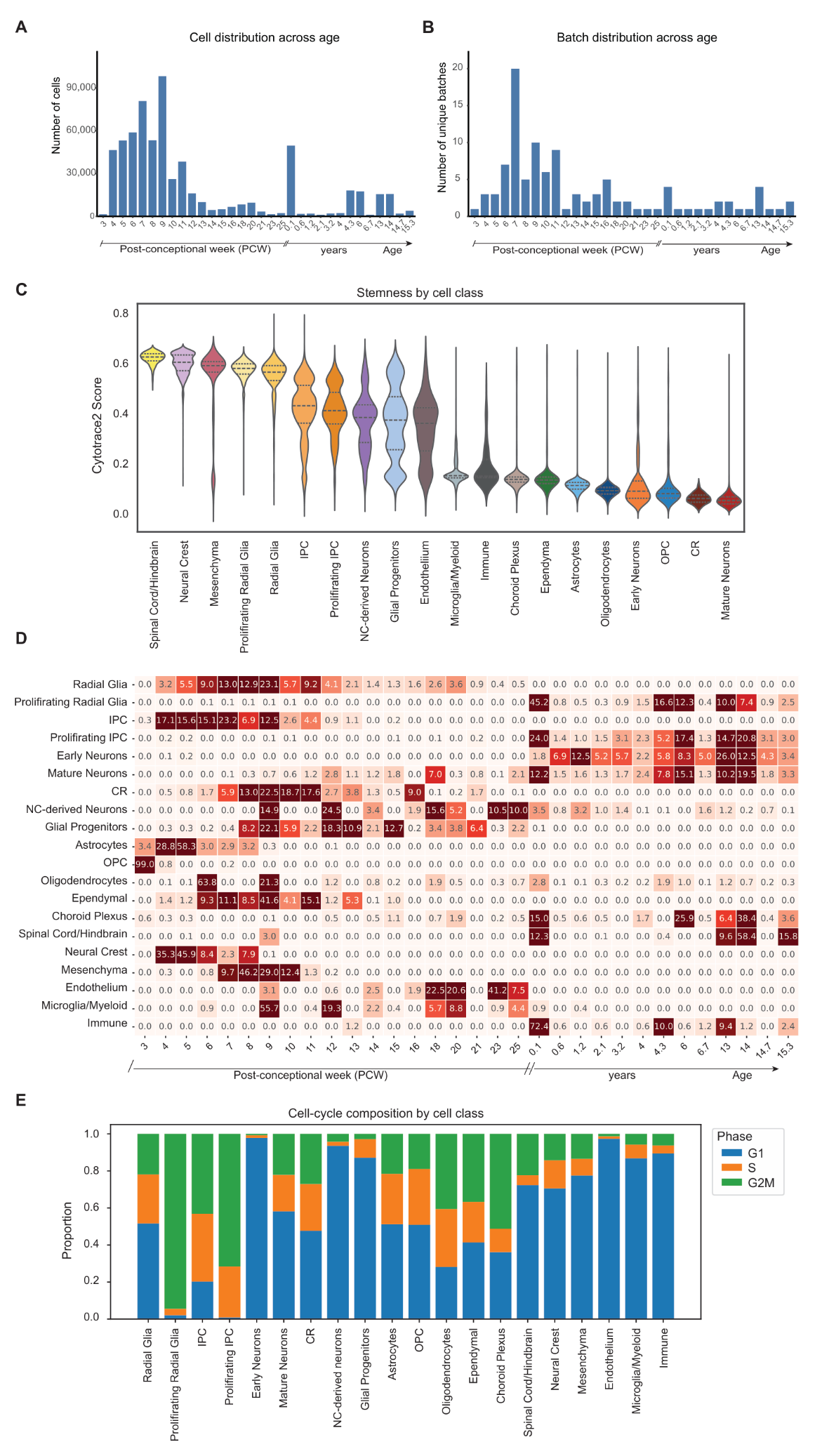

**Figure S1. Developmental composition and cell-state characteristics of HuDeHA.**

**(A)** Distribution of cells across metadata-defined developmental ages in HuDeHA. **(B)** Number of unique batches contributing to each developmental age in HuDeHA.
**(C)** Violin plots of CytoTRACE2 potency scores across annotated cell classes. Higher potency scores were observed in early developmental populations, including neural crest, mesenchymal, and radial glial cells, whereas lower scores characterized more differentiated populations, including astrocytes, oligodendrocytes, and mature neurons.
**(D)** Heatmap showing the distribution of annotated cell classes across metadata-provided developmental ages. Early developmental populations were enriched during embryonic and prenatal stages, whereas later-arising cell classes became more prominent postnatally.
**(E)** Cell-cycle phase distribution across cell classes. Proliferative populations were enriched for S and G2/M phases, whereas differentiated populations were predominantly assigned to G1.

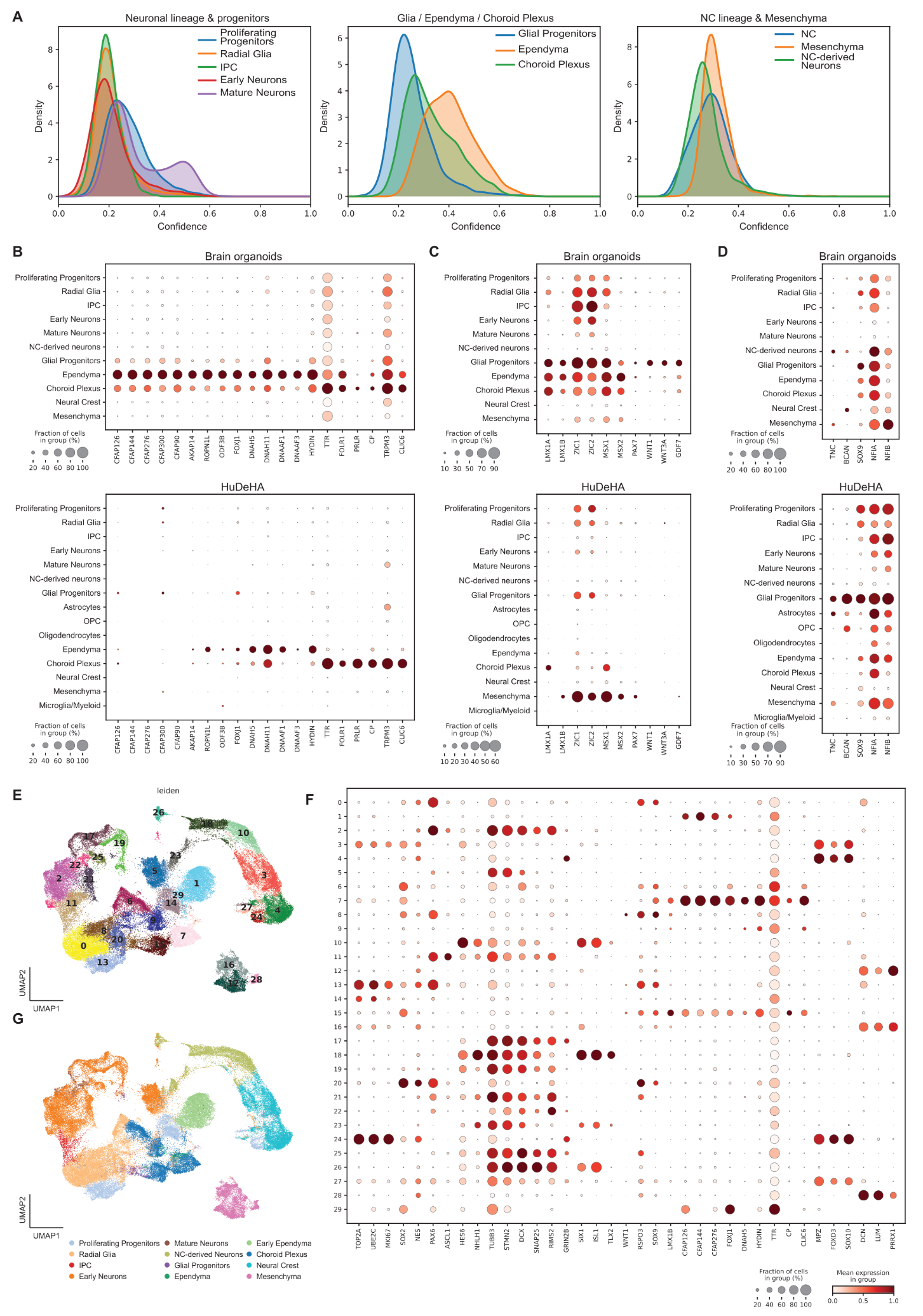

Figure S2. Manual refinement of brain organoid annotations and comparison against HuDeHA.

**(A)** Distribution of scPoli prediction uncertainty across the initially assigned organoid cell classes, grouped into three broad lineage categories. Most cell classes showed relatively low uncertainty, with median values around 0.2. Mature neurons and choroid plexus included subsets of cells with higher uncertainty values of approximately 0.4–0.6, whereas the ependymal population showed a broader overall shift toward higher uncertainty, with a median near 0.4.

**(B)** Dot plot of canonical choroid plexus and ependymal markers in HuDeHA and brain organoids. Canonical mature choroid plexus and ependymal markers are weakly expressed or absent in organoids, whereas CFAP-family genes are enriched, supporting the presence of an organoid-specific ciliated/ependymal-like population.

**(C)** Dot plot showing enrichment of roof plate- and border-associated markers, supporting the presence of a roof plate-like transcriptional program in the organoid population initially annotated as glial progenitors.

**(D)** Dot plot of canonical in vivo glial progenitor markers, showing weak or absent expression in the organoid “glial progenitor” class.
(E) Alternative scVI-based embedding of organoid cells, colored by Leiden clusters, used for manual annotation refinement. Clustering identified three proliferative clusters that were incorporated into the final annotation, as well as a CFAP-enriched cluster annotated as early ependymal cells.
(F) Dot plot of marker genes across Leiden clusters from the refined scVI embedding.
(G) UMAP of organoid cells colored by the final manually refined annotations.

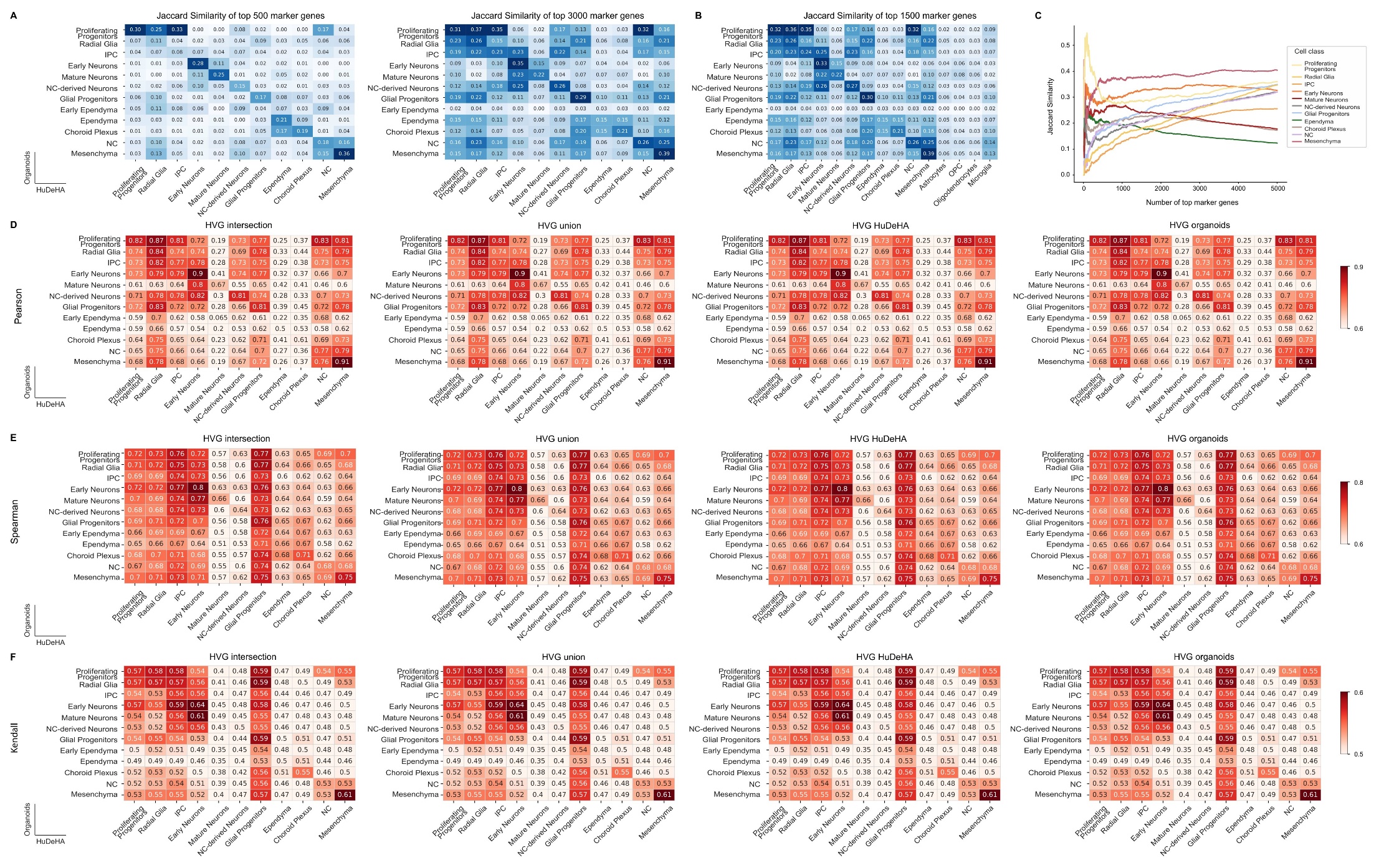

Figure S3. Jaccard and transcriptome-similarity analysis between HuDeHA and brain organoid cell classes.
(A) Jaccard similarity curves comparing matched HuDeHA and brain organoid cell classes using increasing numbers of differentially expressed genes (500, 3000).
(B) Jaccard similarity heatmap comparing all HuDeHA and brain organoid cell classes using the top 1,500 differentially expressed genes. Cell classes not identified in organoids show low similarity to organoid populations.
(C) Jaccard similarity curves showing the effect of the number of genes used for each matched cell class. For most classes, similarity reaches a plateau at approximately 3,000 genes, while proliferating progenitors show the strongest peak at smaller gene numbers.
**(D–F**) Pearson correlation **(D)**, Spearman rank correlation (**E**), and Kendall rank correlation (**F**) between matched HuDeHA and brain organoid cell classes, calculated using four highly variable gene (HVG) sets: the HVG intersection, HVG union, brain organoid HVGs, and HuDeHA HVGs.

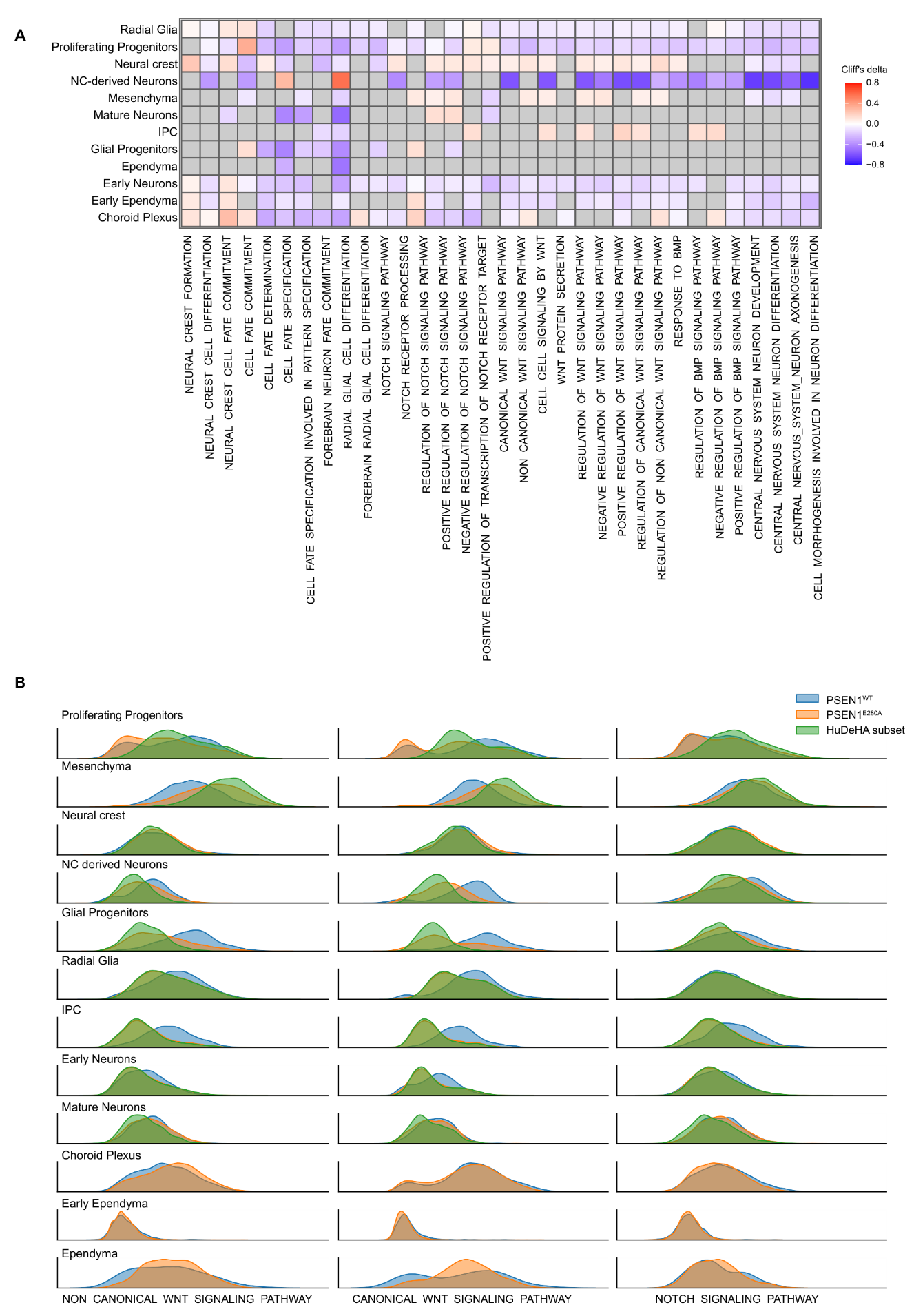

Figure S4. Pathway differences between PSEN1^E280A^ and PSEN1^WT^ brain organoids across cell classes.
(A) Heatmap showing pathway activity differences between PSEN1^E280A^ and PSEN1^WT^ organoids across annotated cell classes. Pathways include WNT, BMP, and Notch signaling, Alzheimer’s disease-related pathways, and neurogenesis/gliogenesis-associated programs. Grey indicates non-significant differences.
(B) Pathway activity histograms comparing PSEN1^E280A^ organoids, PSEN1^WT^ organoids, and age-matched HuDeHA cells for non-canonical WNT signaling, canonical WNT signaling, and Notch signaling.

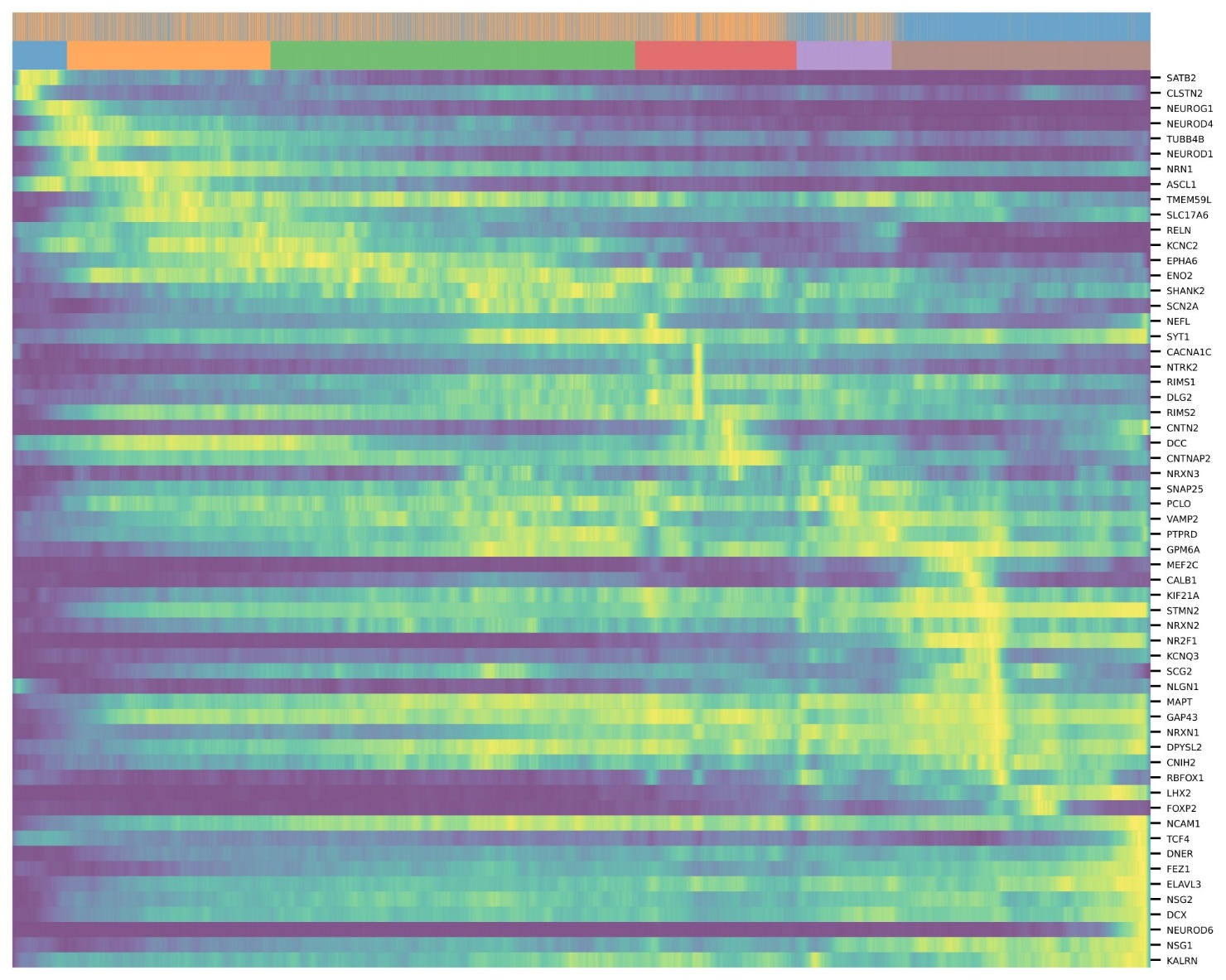

Figure S5. **Dynamic expression of neuronal differentiation genes across organoid pseudotime.**

Heatmap showing genes with the strongest expression changes along the neuronal pseudotime trajectory in brain organoids. Cells are ordered by inferred pseudotime, and genes are shown by scaled expression. The top annotation indicates pseudotime-associated cell-state progression across the neuronal lineage. Genes displayed include markers of early neurogenesis, neuronal differentiation, and neuronal maturation.

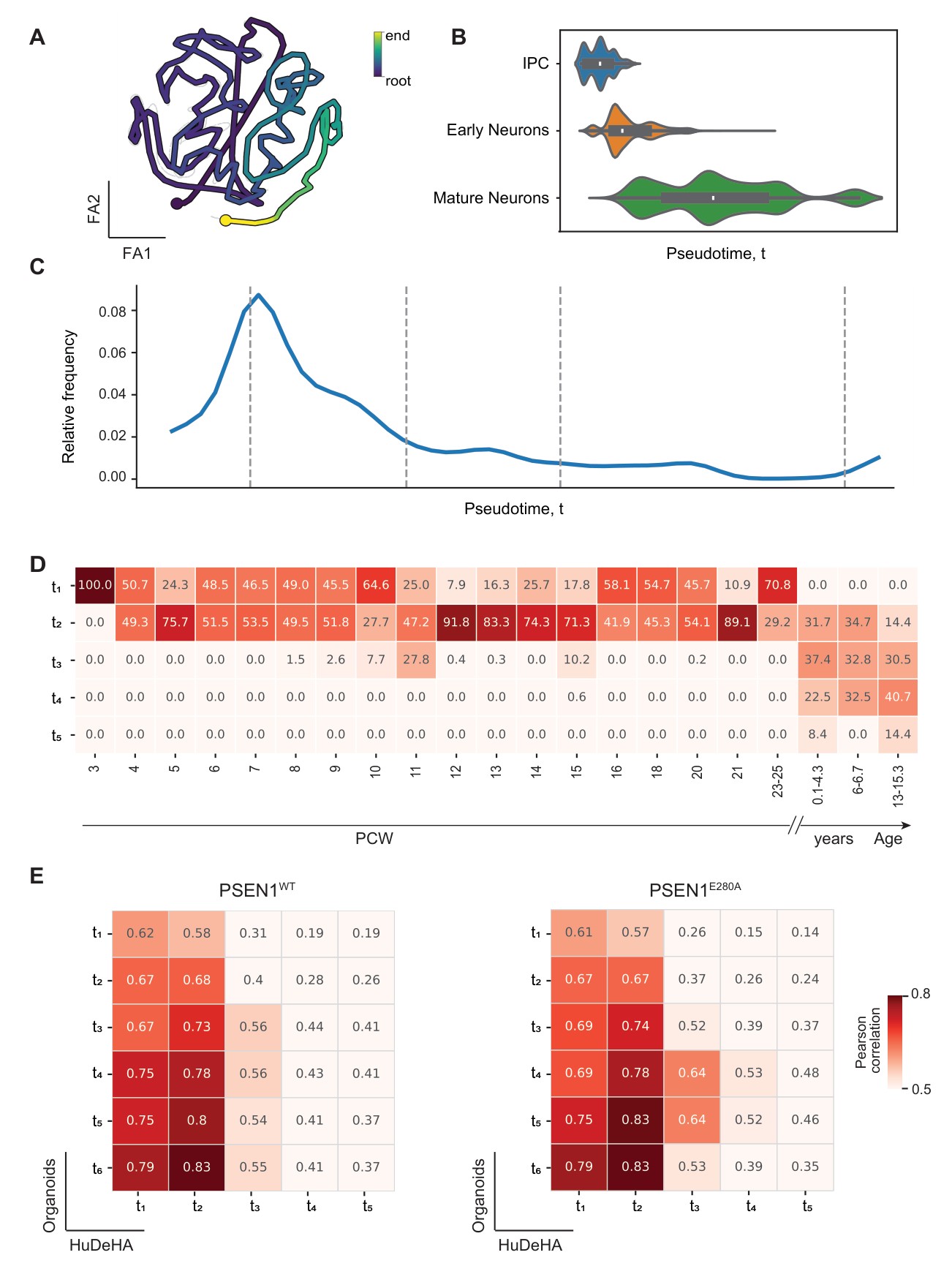

Figure 6. **HuDeHA neuronal pseudotime recapitulates developmental age and enables in vivo–in vitro trajectory comparison.**

**(A)** Reconstructed neuronal lineage trajectory in HuDeHA with cells color-coded by inferred scFates pseudotime, representing a continuous developmental gradient.

**(B)** Violin plots showing the distribution of annotated cell types (IPCs, early neurons, and mature neurons) along the pseudotime axis, confirming that the inferred trajectory aligns with established biological stages.

**(C)** Proportional distribution of HuDeHA neuronal lineage cells across pseudotime, representing in vivo developmental dynamics. Dashed lines indicate custom pseudotime-bin boundaries defined by major inflection points in cell abundance. The absence of strong cell enrichment at the end of the HuDeHA trajectory suggests that the late-pseudotime shift observed in PSEN1^E280A^ organoids reflects a genotype-associated effect rather than an artifact of the reference trajectory structure.

**(D)** Heatmap showing the percentage of HuDeHA cells mapped to metadata-provided developmental timepoints across pseudotime bins. The alignment confirms that earlier pseudotime bins correspond to earlier biological developmental ages.

**(E)** In vivo–in vitro transcriptomic similarity shown by heatmaps displaying Pearson correlations between HuDeHA and brain organoid pseudotime bins, computed using shared highly variable genes across PSEN1 genotypes. The correlation pattern indicates that PSEN1^E280A^ organoids are shifted toward more mature HuDeHA time bins, supporting accelerated neuronal maturation in the mutant condition.

**Supplementary Tables
Table S1** – Batch-level metadata for all datasets included in HuDeHA.
**Table S2** – Top 30 upregulated and downregulated differentially expressed gene between PSEN1^E280A^ and PSEN1^WT^ brain organoids for each cell class.
**Table S3** – Top 30 marker genes defining each annotated cell class in HuDeHA.
**Table S4** – Top 30 marker genes defining each annotated cell class in iPSC-derived brain organoids.
